# Distinct timescales for the neuronal encoding of vocal signals in a high-order auditory area

**DOI:** 10.1101/2021.06.28.450187

**Authors:** Aurore Cazala, Catherine Del Negro, Nicolas Giret

## Abstract

The ability of the auditory system to selectively recognize natural sound categories with a tolerance to variations within categories is thought to be crucial for vocal communication. Subtle variations, however, may have functional roles. To date, how the coding of the balance between tolerance and sensitivity to variations in acoustic signals is performed at the neuronal level requires further studies. We investigated whether neurons of a high-order auditory area in a songbird species, the zebra finch, are sensitive to natural variations in vocal signals by recording responses to repeated exposure to similar and variant sound sequences. We took advantage of the intensive repetition of the male songs which subtly vary from rendition to rendition. In both anesthetized and awake birds, responses based on firing rate during sequence presentation did not show any clear sensitivity to these variations, unlike the temporal reliability of responses based on a 10 milliseconds resolution that depended on whether variant or similar sequences were broadcasted and the context of presentation. Results therefore suggest that auditory processing operates on distinct timescales, a short one to detect variations in individual’s vocal signals, longer ones that allow tolerance in vocal signal structure and the encoding of the global context.

## Introduction

Vocal communication signals may provide rich information through both their acoustic structure and subtle variations in their acoustic features^1,2^. A given word spoken by various people convey information about its meaning through an invariant acoustic structure among uttered signals. It may also provide information about the gender, the emotional state and the individual identity of the emitter through fine variations in temporal and acoustic features of uttered signals across individuals. Vocal communication is therefore a computational challenge, requiring the auditory system to selectively extract invariant information with a tolerance to variations for categorization but with sensitivity to variations that potentially provide supplementary information^3^. Within this framework, how the balance between tolerance and sensitivity to subtle variations in acoustic signals is encoded at the neuronal level within the auditory system still require further investigations^4–6^.

Songbirds offer a powerful model to explore neural coding principles underlying this balance. Birdsong is a complex multiple cues signal that is pertinent to species identity and exhibits subtle variations that may carry information such as group or individual identity, emotional or motivational state or physical conditions^7,8^. Among songbird species, the zebra finch is very well suited for investigating how subtle variations encompassed within highly similar communication sounds are encoded within the auditory system. The male zebra finch typically produces a single individual-specific stereotyped song motif that includes several distinctive sound elements, called syllables, that are always produced in the same order^9^. In spite of high stereotypy in their acoustic structure, motifs vary from rendition to rendition with a degree of variations carrying information about the social context, *i.e*. the presence or absence of females^10^. Also, a recent study provides evidence that subtle variations can be perceived by zebra finches^11^. Male zebra finches intensively repeat their song everyday while repetition of the same stimulus is well-known to elicit habituation in behavioral and neural responses raising the question whether variations could have an impact on these changes in responses.

In songbirds, the processing of complex behaviorally relevant acoustic signals, including calls and songs, involves an auditory area analogous to secondary auditory cortex in mammals, the caudomedial nidopallium (NCM), that is a good candidate for investigating how the balance between tolerance and sensitivity to subtle variations in acoustic signals is encoded^3^. Neurons in this auditory area display a clear preference for natural over artificial sounds. Regarding conspecific vocal signals, they may exhibit invariant responses to call categories^12,13^. In spite of this tolerance to variations in vocal signals, neurons in NCM also support recognition of familiar vocalizations that only differ in fine acoustic detail among their categories^14–16^. Neurons in NCM also display stimulus-specific adaptation during which the repeated exposure to a given auditory stimulus induces a decrease in responses and the exposure to a novel stimulus or to the same stimulus with a different order of the sound elements resets responses^15,17–20^. To date, this phenomenon, interpreted as reflecting memory formation, was reported only in experiments in which the exactly same sound stimuli were repeatedly presented. However, in the wild, individuals are never exposed to similar vocal signals as fine natural variations in acoustic features always occur across renditions, raising the question whether these variations might affect neuronal responses in NCM and their time course. Based on extracellular recordings in both anesthetized and awake zebra finches, we show a clear impact of these subtle variations on neuronal responses driven by sequences of song elements that either varied in acoustic details or remained the same across renditions. This impact was observed in spike timing and at a short temporal resolution reflecting a temporal integration of acoustic features across different time scales.

## Results

To explore the neuronal sensitivity to subtle acoustic variations across renditions of vocal signals in a high-order auditory area, we performed extracellular recordings of NCM neurons in awake zebra finches (n=4 birds) while playing back sequences built from individual’s song syllables. These sequences were arranged in two different sound series, the ABAB-Same and the ABAB-Var series, both consisting of two song syllables, called A and B, repeated twice alternatively to form an ABAB sequence. The ABAB-Same series were built from 60 repetitions of a single ABAB sequence while the ABAB-Var series from 60 natural variants of a given ABAB sequence (Fig. 1a-c). The similarity in fine acoustic structure of A or B syllables from one sequence variant to another was evaluated using the percent accuracy score in Sound Analysis Pro 2011^21^. Renditions of A or B syllables from one variant to another in ABAB-Var sequences were, on average, 83.2% and 81.9% similar, respectively, while, in comparison A and B syllables within a given sequence were significantly less similar, on average 73.5% in ABAB-Same sequences and 68.8% in ABAB-Var sequences (t-tests, *p* < 0.001; Fig. 1c).

**Figure 1:**
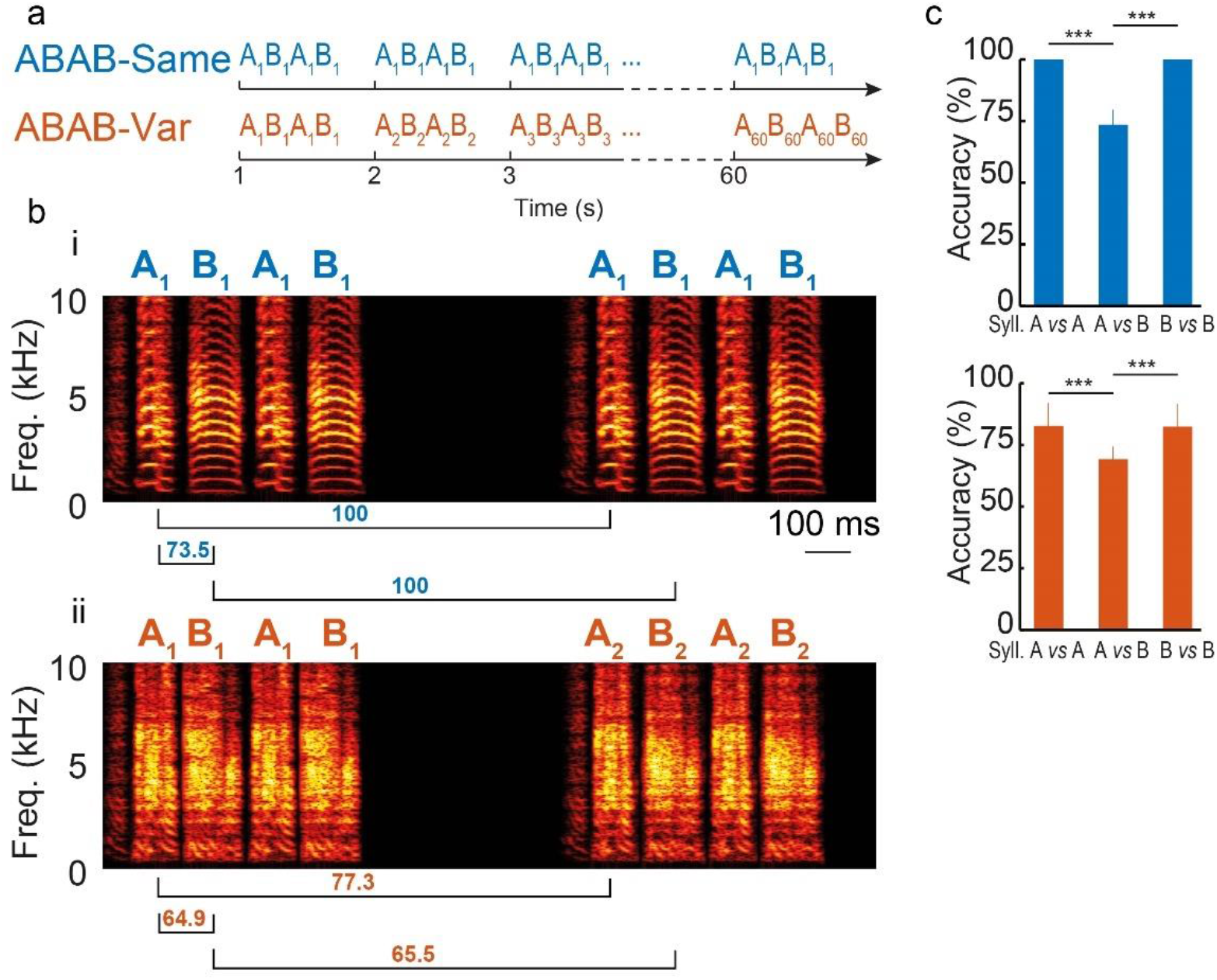
A single sequence or sequences with natural variations found in individual’s songs were used to build two series types: ABAB-Same and ABAB-Var series. a) Schematic diagram of the structure of ABAB-Same (top) and ABAB-Var (bottom) series. A and B depict two syllable types used to form ABAB sequences. The ABAB-Same series consisted of 60 repetitions of a single ABAB sequence while the ABAB-Var series consisted of 60 distinct renditions of a given ABAB sequence. These renditions called sequence “variants” were labelled as A_n_B_n_A_n_B_n_ (n varying from 1 to 60). A_n_ and B_n_ were distinct exemplars of a single syllable type that were extracted from the song’s repertoire of a given individual. Each sequence was presented at a rate of one per second. b) Example spectrograms of two consecutive sequences within an ABAB-Same (i, no variants) and ABAB-Var (ii, variants) series. Note the subtle changes between A_1_B_1_A_1_B_1_ and A_2_B_2_A_2_B_2_ sequences of the ABAB-Var serie (e.g. power at ~5kHz on syllable B. Underneath each spectrogram are the accuracy scores (%) computed with SAP 2011 (see main text for further details) between A and B syllables across the two successive example renditions of the ABAB-Same and ABAB-Var sequences. c) Mean (+/- STD) of the accuracy scores computed between A and B syllables across the 60 renditions of all the ABAB-Same (top) and ABAB-Var (bottom) sequences. *** *p* < 0.001.

### No effect of acoustic variations on response strength in awake birds

To assess auditory responses to playbacks of ABAB-Same and ABAB-Var series in awake birds, we performed three (range: 2-5) recording sessions (3.6 electrodes per recording session, range 2-7) per bird, with 4.5 days (range 1-9) between two successive recording sessions. We analyzed the spiking activity of 56 recording sites, located from the dorsorostral portion (maximal depth 2000 μm) to the dorsocaudal portion^20^. They were driven by the playback of the ABAB-Same and ABAB-Var sequences, as illustrated by the example unit on Fig. 2a-b.

**Figure 2:**
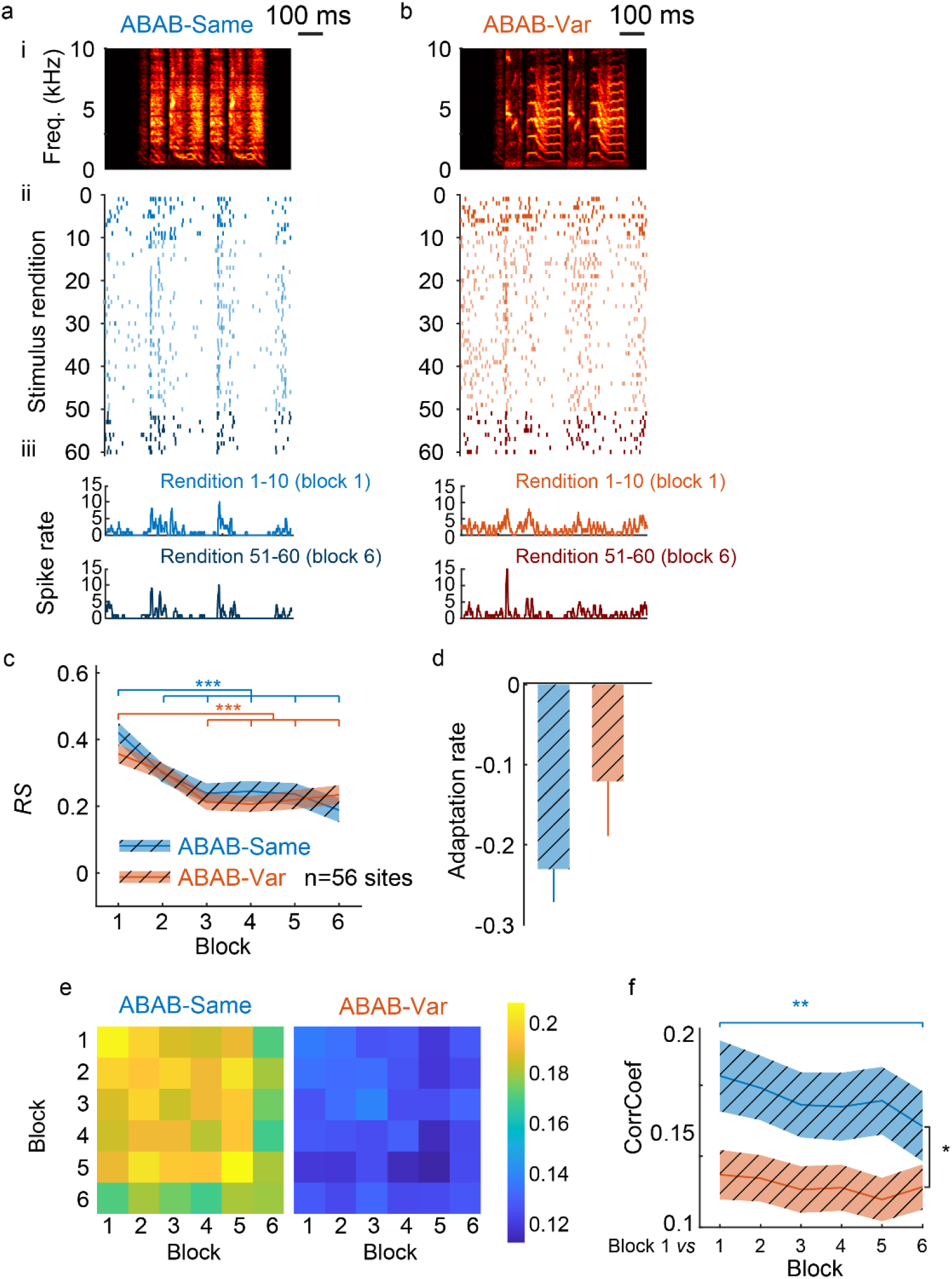
Auditory responses to 60 repetitions of a single sequence (ABAB-Same series) and to 60 sequence variants (ABAB-Var series) in awake birds. Responses of a representative unit to the ABAB-Same (a) and the ABAB-Var (b) series used as auditory stimuli. Neuronal responses are shown as raster plots (60 iterations) and peristimulus time histograms (bottom; 10 ms bin width; for the 10 first and the 10 last trials) that are time-aligned with sequence spectrograms (top: the sequence repeated 60 times for the ABAB-Same example series and one sequence variant for the ABAB-Var example series). (c) Modulation of responses over the 6 successive blocks of ten trials (blocks for the ABAB-Var series include 10 variants of the auditory sequence). The *RS* values estimated the strength of the responses driven by the series used as auditory stimulus. Thick line indicates mean responses for the population of recording sites (n=56). Hatched area represents SEM. (d) Adaptation rate (mean ± SEM) of responses computed over the 10 first trials did not significantly differ between the two series. (e) Reliability of spike trains illustrated by heatmaps (right: ABAB-Same series; left: ABAB-Var series). Spike trains reliability, quantified by the CorrCoef index, was lower when sequence variants were presented. Blue color indicates low CorrCoef values. (f) At the population level, differences in spike-timing reliability and in its time course between the two series. CorrCoef values were computed from spike trains evoked by the first ten trials and those evoked by the ten trials of the six blocks (block 1 to 6). CorrCoef computed for block 1 *vs* 1 is not equal to 1 because it is computed on each iteration (e.g. iteration m *vs* iteration n, with *m* and *n* ranging from 1 to 10). Significant difference: * p<0.05, ** *p* < 0.01, *** p< 0.001 (see main text for statistics details).

To examine whether the time course of auditory responses differed between the ABAB-Same and the ABAB-Var series, we performed a repeated-measures (RM) ANOVA on the response strength (*RS*), computed from firing rates averaged over the entire sequence duration, using a linear mixed-effect model with sequence type and block repetition as cofactors and units as a random factor (Fig. 2c). We used the term “block” because data were averaged over 10 trials, but all trials were delivered at the same frequency, one trial per second. Results indicated that response strength did not differ between ABAB-Same and ABAB-Var series (sequence type factor; F_1, 564_ = 0.03, *p* = 0.85). Numerous studies have reported a stimulus-specific adaptation of auditory responses in NCM when the playbacks of conspecific vocalizations are repeated^15,17,19,20,22^. The RM ANOVA revealed an effect of block repetition factor on *RS* values (F_5, 564_ = 30.38, *p* < 0.0001) with a decrease in the strength of responses to both series (post-hoc tests: ABAB-Same: block 1 *vs*. block 2 to 6 all *p* < 0.001; ABAB-Var: block 1 *vs*. block 3 to 6, all *p* < 0.001). Statistical analysis also revealed a significant interaction between block repetition and series type factors (F_5, 564_ = 2.26, *p* = 0.047) suggesting that the time course of auditory responses over the 60 renditions of ABAB sequences depended on whether acoustic features of syllables varied or not. Responses changed dramatically over the first stimulus presentations^22^. Here, NCM neurons displayed a significant decrease in their activity from the first block to the second one when ABAB-Same series were played back, leading us to examine whether responses of NCM neurons adapted more rapidly to the ABAB-Same series than to the ABAB-Var ones. We computed the adaptation rate for both sequences by extracting the slope of the linear regression over the 10 first stimulus renditions for each unit, as in several previous studies^17,23–25^. Although the average adaptation rate was higher for ABAB-Same than ABAB-Var sequences (Fig. 2d), it did not significantly differ (t_1, 55_ = 1.18, *p* = 0.24). These results therefore indicate no clear effect of rendition-to-rendition acoustic variations in syllable features on the time course of neuronal responses.

### Impact of acoustic variations in spike-timing reliability in awake birds

We analyzed the temporal pattern of auditory responses by computing the trial-to-trial reliability coefficient, the CorrCoef. High CorrCoef values indicate a high spike train reliability across trials while low CorrCoef values mean great variations in temporal patterns of spike trains. This coefficient was calculated using responses over 20 presentations, the ten presentations of sequence stimuli of the a given block and those of each of the 6 blocks. Results indicated that CorrCoef values varied between [-0.07 and 0.69] with an average of 0.13, which is in the range usually reported for cortical^26–28^ and NCM neurons^20^.

Analyses of CorrCoef values revealed an impact of series type and block repetition (linear mixed effect model, RM ANOVA; series type factor; F_1, 110_ = 4.73, *p* = 0.032; block repetition factor, F_5, 550_ = 3.62, *p* = 0.003). The trial-to-trial spike-timing reliability was significantly lower when ABAB-Var series were played back (Fig 2e) suggesting greater variations in spike-timing of responses when sequences consisted of ABAB variants than when the same sequence was repeatedly played back. Post-hoc tests focused on comparisons between the first block and the other ones revealed that the trial-to-trial reliability of spike trains was modulated by the repetition of the same ABAB sequence, CorrCoef values significantly decreasing with sequence renditions (Fig 2f; block 1/block 1 *vs*. block1/block 6; *p* = 0.0027). In contrast, the trial-to-trial reliability of spike trains evoked by variants in ABAB-Var series remained lower and stable (p > 0.68; see heatmaps on Fig. 2e). The accuracy of spike timing continued to vary considerably throughout the exposure to the variants.

### Auditory responses to variant and similar sequences in anesthetized birds

Extracellular recordings in NCM were also performed in seven isoflurane-anesthetized adult males. Only well-isolated responsive single units (n=82) were selected (example unit on Fig. 3a-b). These single units were from the dorsorostral portion (maximal depth 2000 μm) to the dorsocaudal portion and they were driven by the playback of the ABAB-Same and the ABAB-Var series.

**Figure 3:**
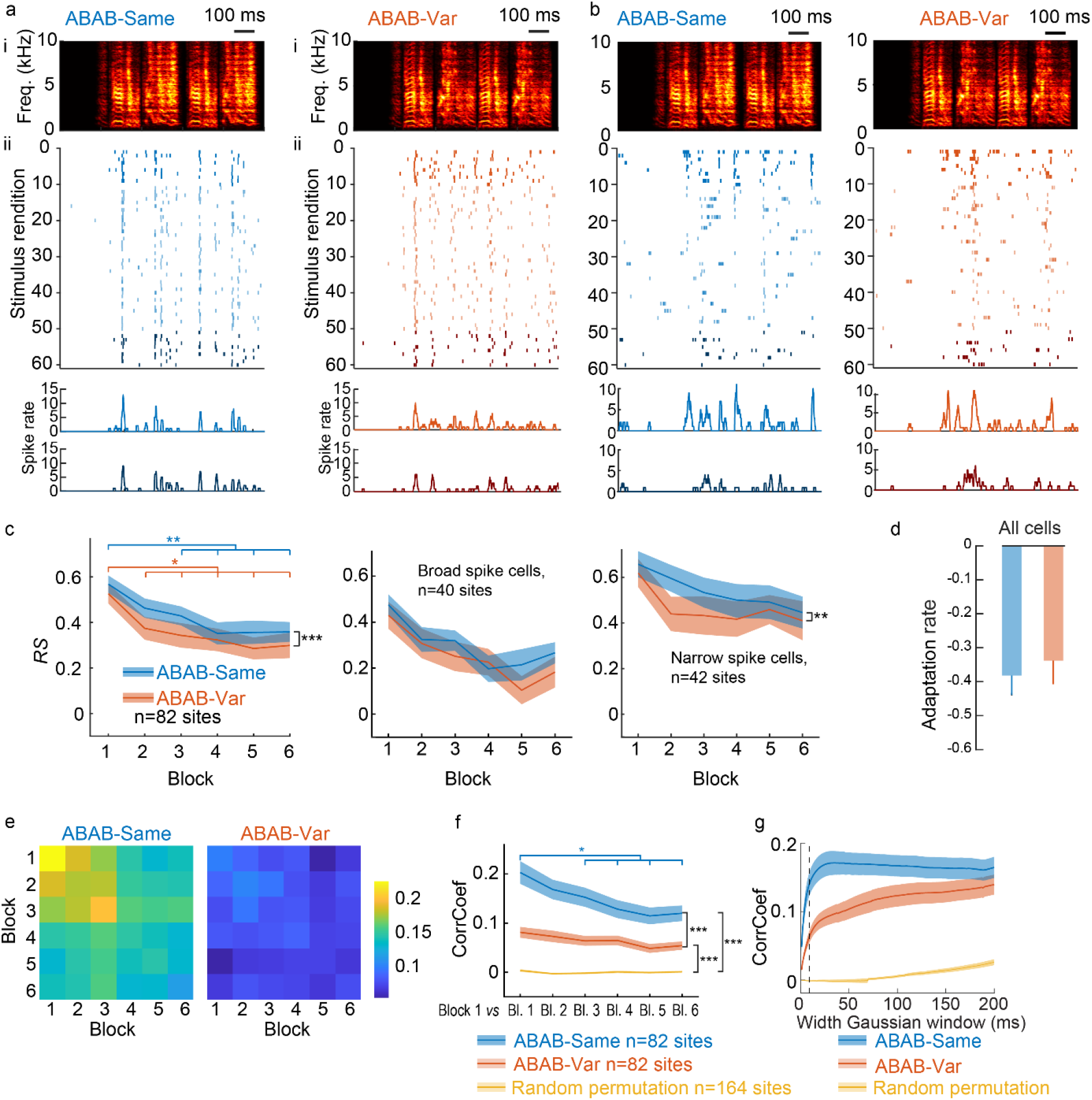
Auditory responses in anesthetized birds. From rendition to rendition, spike timing greatly changed when sequence variants were played back. No such changes were observed when the same sequence was repeated (ABAB-Same series). Neuronal responses of a representative single unit to playback of one ABAB-Same (a) and one ABAB-Var (b) series are shown as raster plots (60 iterations) and peristimulus time histograms (bottom; 10 ms bin width; for the 10 first and the 10 last trials) that are time-aligned with sequence spectrograms (top: the sequence repeated 60 times for the ABAB-Same example series and one sequence variant for the ABAB-Var example series). c) ABAB-Same series evoked higher responses (*RS* values) than ABAB-Same series at the population level (left) and for the sub-population of narrow spike cells (right), but not for broad spike cells (middle). Thick line indicates mean values and shaded area represents SEM. d) Response strength differed, but similarly changed with repeated exposure to sequences, as indicated by the adaptation rate computed over the first ten stimuli presentations (mean ± SEM). e) As observed in awake birds, spike train reliability differed between the two series, with a higher spike timing accuracy when the same sequence (ABAB-Same) was repeated. Heatmaps from CorrCoef values computed per block of 10 stimuli renditions. f) Corrcoef (mean ± SEM) changed with stimulus exposure when the same sequence was repeated while it remained similar when sequence variants were played back. Corrcoef values were higher than those of spike trains in which spike timing was randomly permutated. g) Varying the Gaussian window width used to compute the convolution of spike trains from 1 to 200 ms affects CorrCoef values. In the present study, a 10 ms Gaussian window width to compute CorrCoef values (vertical dashed line) and Corrcoef values differed between the two series. No difference between ABAB-Same and ABAB-Var was observed when the time window exceeds 98 ms. CorrCoef values were also computed on spike trains after a random permutation of the spike timing. Thick line indicates mean values; shaded area represents SEM. Significant difference: \**p* <0.05, ** *p* < 0.01, *** *p* < 0.001.

The RM ANOVA performed on *RS* values revealed that they differed between ABAB-Same and ABAB-Var series over the six blocks (series type factor: F_1, 1055_ = 12.87, *p* = 0.0003). However, auditory responses did not differ when comparisons were focused on each block (post-hoc tests; all *p* > 0.64). As in awake birds, neuronal responses showed the well-described adaptation across stimulus presentations (block repetition factor: F_5, 1055_ = 13.02,*p* < 0.0001). Both series induced a significant decrease over block repetitions (ABAB-Same series: block 1 *vs*. block 3,4, 5 and 6, all *p* <0.01; ABAB-Var series: block 1 *vs*. block, p <0.0021, block 1 *vs*. block 3, 4, 5 and 6, p<0,01) with no difference in adaptation rate over the ten first trials (F_1, 81_ = 0.74, *p* = 0.46). Therefore, subtle variations in acoustic features of syllables in ABAB-Var series had no clear impact on responses on the basis of firing rate measures.

Two cell types can be distinguished in NCM^3,20,29–31^. Responsive NCM neurons were split into two populations according to the peak-to-peak width of their action potential: neurons with broad spikes (≥0.3 ms; *n* = 40, width = 0.49+/-0.10 ms) and neurons with narrow spikes (<0.3 ms; *n* = 42, width = 0.27+/-0.07 ms). The RM ANOVA performed on *RS* values according to the block repetition revealed a significant decrease in response strength of both cell types (broad-spike cells, linear-mixed effect: F_5,428_ = 9.29, *p* < 0.0001; narrow-spikes cells, linear-mixed effect: F_5,448_ = 5.01, *p* < 0.0003) and a significant series type effect for narrow-spikes cells (broad-spike cells, series type factor: F_1,428_ = 3.10, *p* = 0.08; narrow-spikes cells, series type factor: F_1, 448_ = 7.72, *p* < 0.006), but no significant interaction between the two factors for both cell types (broad-spike cells, F_5,428_ = 0.53, *p* = 0.75; narrow-spikes cells, F_5,448_ = 0.55, *p* = 0.73). When the analysis was focused on the first ten renditions of the first block, both cell types did not show any effect of natural variations on adaptation rate (broad-spike cells, paired t-test: t_38_ = 1.39, *p* = 0.17; narrow-spike cells, paired t-test: t_40_ = 0.26, *p* = 0.79; note that for both cell types, one unit was removed because it did not spike during the first trial).

### Impact of acoustic variations in spike-timing reliability in anesthetized birds

We also evaluated the spike timing reliability across blocks of sequence presentations by computing the CorrCoef. Most of the results are consistent with those obtained in awake birds. As illustrated by Fig. 3e-f, CorrCoef values were higher for ABAB-Same than for ABAB-Var series (series type, F_1, 891_ = 199.32, *p* < 0.0001; Fig. 3f) suggesting that spike trains were more reliable across the iterations of the same sequence than across the renditions of variants. Importantly, CorrCoef values of spike trains evoked by variants were significantly higher than CorrCoef values of spike trains in which inter-spike times were randomly distributed (RM ANOVA, series type: F_2,1869_ = 501.09, *p* < 0.0001; post-hoc test: ABAB-Same *vs* Random permutation, *p* < 0.0001; ABAB-Var *vs* Random permutation, *p* < 0.0001; yellow line in Fig. 3f). This points out a certain degree of trial-to-trial reliability in spike trains evoked by variants.

Spike train reliability gradually decreases reaching a significant decrease from the third block, when the same sequence within ABAB-Same series was repeatedly played back (block repetition factor: F_5, 891_ = 10.52,*p* < 0.001; block 1 *vs* block 2: *p* = 0.31; block 1 *vs* block 3: *p* = 0.024; block 1 *vs* block 4 to 6: multiple *p* < 0.001; Fig. 3d). Such decrease in CorrCoef values was not observed when ABAB-Var series were used as stimuli (multiple *p* > 0.13; Fig. 3f). Therefore, as in awake birds, the temporal reliability of spike trains remained stable, showing no clear effect of the repeated exposure to sequence variants.

Here, CorrCoef were computed after applying a convolution on spike trains with a 10 ms Gaussian window width, a time resolution considered as optimal for discrimination of conspecific songs in auditory structures^28,32,33^. Using this 10 ms time resolution, CorrCoef results showed a sensitivity to natural variations in individual’s vocal signals that failed to show results based on firing rates averaged over the several hundreds of milliseconds of the whole sequence duration. To bridge the gap between the two timescales, 10 milliseconds *vs*. several hundreds of milliseconds, we computed CorrCoef varying the width of the Gaussian window from 1 to 200 milliseconds. Importantly, as the width of the Gaussian window increases, spike trains are more and more smoothed and so, the trial-to-trial reliability of spike trains becomes increasingly based on firing rate rather than on spike timing accuracy. Our aim was to determine the time resolution where CorrCoef values did no longer differ between the two series. As shown in Fig. 3g, while CorrCoef values reached a plateau with a Gaussian window width at about 10 ms when ABAB-Same series were played back (Fig. 3g), CorrCoef values remained lower up to 170 ms for spike trains evoked by variants, both CorrCoef values being always much higher that after a random permutation of the spike times. As the time scale was increasing, the difference in CorrCoef values between ABAB-Same and -Var was decreasing with no significant difference when the width of the Gaussian window was higher than 98 ms (linear mixed-effect models at each time point). This suggests that sensitivity to natural subtle variations in acoustic features across variant renditions requires a short time scale (< 100 ms) that fits within the duration range of syllables [63.5 – 203.6 ms] used to form sequences in the present study.

### No relationships between responses and variations in auditory stimuli

Variations in temporal and acoustic syllable features across variant renditions offered us the possibility to examine to what extent the trial-to-trial variability in spike train accuracy relied on the degree of variations in syllable features across renditions. To address this issue, we examined to what extent variations in syllable length contributed to the reliability of spike trains by performing a linear time warping that allows aligning all spike-trains evoked by individual A and B syllables of ABAB-Var series on a common time axis (see Methods). This method reduces variability in the alignment of syllables onset and offset. A paired t-test on CorrCoef values obtained after comparing spike trains between blocks revealed that time warping significantly changed CorrCoef values (t_20_ = −2.60, *p* =0.017). However, this change was small, CorrCoef values being marginally changed after time warping (before: 0.081 ± −0.01 *vs* after: 0.083 ± 0.01, mean ± STD) and CorrCoef values remained significantly different between ABAB-Same and -Var series after time warping (mean ± STD = 0.16 ± −0.026; t_20_ = −17.6, *p* <0.0001). Variations in syllable length therefore explained only a small part of the lower reliability of spike trains evoked by ABAB-Var series. We then assessed whether the more two variants were acoustically different, the lower the reliability of spike trains evoked by these two variants. Similarity scores, entropy and pitch differences between the first sequence and the 59 subsequent ones in ABAB-Var series were computed using Sound Analysis Pro^21^. In parallel, we calculated CorrCoef values between the spike train evoked by the first sequence of the ABAB-Var series used as stimulus and those evoked by the 59 others. Similarity score that describes the acoustic similarity of a pair of sound stimuli based on several acoustic parameters confirmed the subtle variations in fine acoustic structure of syllables, this measure (mean ± SD: 96.32% ± 3.60, range: [54-100 %]). Linear regressions based on either similarity scores, entropy or pitch differences and CorrCoef values did not reveal any significant correlations (*p* > 0.15; Fig 4b-d). Thus, results did not show any relationships between trial-to trial reliability of spike trains and the degree of variability in acoustic features across renditions. These results therefore provide additional support for a non-linear processing of acoustic features^20,29,34–36^.

**Figure 4:**
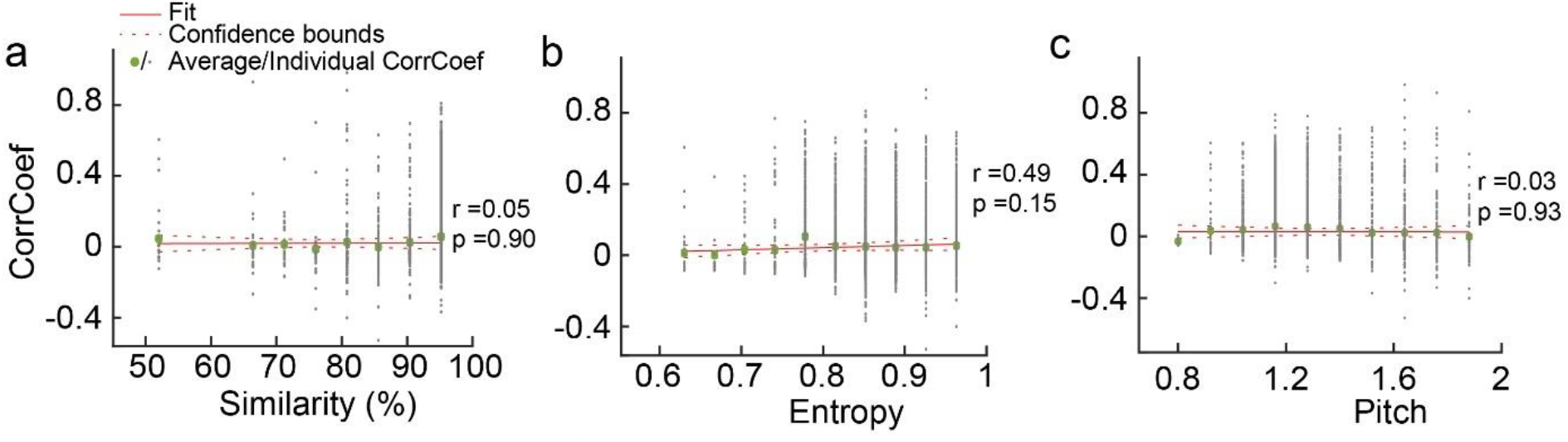
No correlation between variability in acoustic features and spike timing reliability. a) Response strength in both anesthetized (open boxes) and awake birds (dashed boxes) did not depend on the exemplar of series used as stimuli, *i.e*. on the syllable types used to form sequences within the series. Eight ABAB-Var series and seven ABAB-Same series were used as stimuli. Numbers in black and grey below bars indicates how many times the corresponding playback file was used and how many neurons of the overall population of recorded neurons responded to the series type, respectively. Note that the ABAB-Var series labelled as S8 that induced the greatest auditory response was presented only once. b-d) Linear regression between differences in similarity scores (b), entropy (c) and pitch (d) from the first sequence exemplar of the ABAB-Var series and one of the 59 following ones *vs* CorrCoef values, computed from the spike train evoked by the first sequence rendition and one of the 59 following ones, the same as used to quantify acoustic differences. The thick line represents the slope of the regression; Pearson’r and *p* values on each plot; green dot: averaged CorrCoef values.

### Effect of context on the repetition of the AB pair within sequences

Neurons in NCM are sensitive to sequence ordering and context^20,29^. Sequence stimuli used in ABAB-Same and ABAB-Var series were all built from a given pair of AB syllables repeated twice. What differed between ABAB-Same and ABAB-Var series was the context in which ABAB sequences occurred: the same sequence *vs*. various versions of the sequence. We took advantage of the repetition of a given AB pair within sequences and the difference in context between the two series to assess whether the type of context affected responses to the second rendition of AB pair within ABAB sequences. In awake birds, analyses of *RS* values revealed a significant decrease in responses with AB pair repetition within sequences of both series (F_1,172_ = 5.90, *p* < 0.02; Fig. 5a) but with no difference between the two series (F_1,172_ = 0.32, *p* = 0.57) and no significant interaction between the two factors (F_1,172_ = 3.34, *p* = 0.07). Analyses of spike timing accuracy using CorrCoef values also pointed out an impact of AB pair repetition on responses (F_1,172_ = 24.42, *p* < 0.0001; Fig. 5b). Interestingly, the effect of AB pair repetition was observed when ABAB-Same as well as ABAB-Var series were played back (post-hoc tests, *p* < 0.01 and *p* < 0.001, respectively) indicating that, even if CorrCoef values for spike trains evoked by ABAB-Var series were low, they could reveal changes in spike train accuracy. The temporal pattern of discharges was, therefore, impacted by the AB pair repetition in both contexts. However, the trial-by-trial comparisons of spike trains evoked by each of the two AB pairs based on the Pearson correlation coefficient indicated a significant difference between ABAB-Same and -Var series (paired t-test, t_55_ = −2.07, *p* = 0.043; Fig. 5c) with a higher effect of the AB pair repetition on temporal pattern of spike trains in ABAB-Var series. These results therefore provide evidence of an impact of the context on auditory responses in NCM.

**Figure 5:**
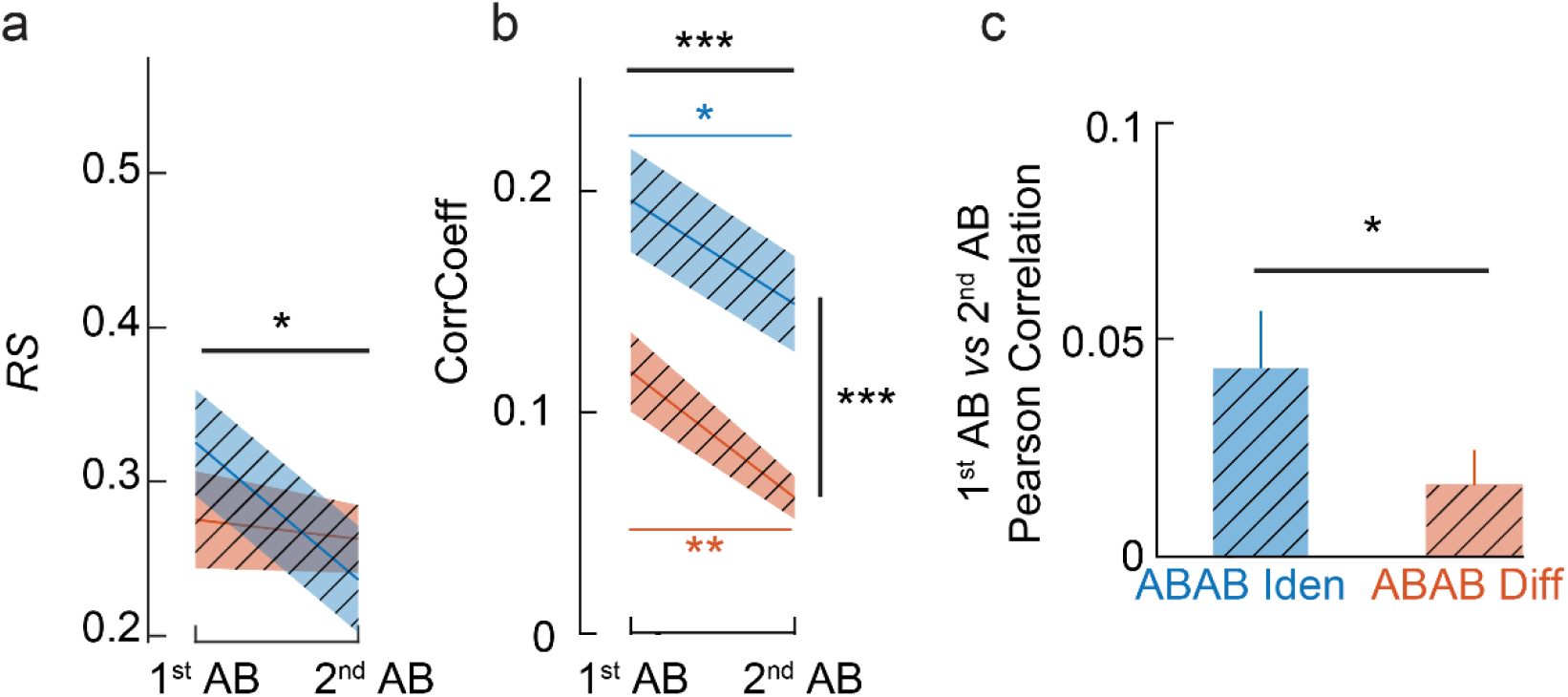
Responses to the two AB pairs that form ABAB sequences reflects sensitivity to the context in awake birds. (a) Strength of responses (*RS* values) changed from the first AB pair to the second one. The exposure to the first pair of syllables AB impacts the responses to the second pair of syllables AB within a stimulus rendition in both anaesthetized and awake birds. Evoked auditory responses (a) and CorrCoef (b) were overall higher for ABAB-Same than for ABAB-Var sequences and were lower for the second pair of syllables AB than for the first pair. Yet, Pearson correlation coefficient measured on each individual spike train between the first and second pair of syllables AB was lower for ABAB-Var than ABAB-Same sequences (c). *, ** and ***, *p* < 0.05, 0.01 and 0.001, respectively (see main text for statistics details).

## Discussion

Across renditions, vocal signals acoustically vary, raising the question whether these variations are detected and play functional roles. Subtle natural variations in fine acoustic structure of song syllables can be behaviorally discriminated by adult zebra finches^11^. Our study provides evidence that these variations are encoded by neurons of a high-level auditory area, as indicated by spike train reliability that differ depending on whether acoustic details vary across iterations.

With regard to the functional role, we aimed at investigating the impact of natural variations on the adaptation of neural responses to a repeated stimulus, that is considered as playing a role in auditory memory formation through the binding of auditory objects, a crucial processing of the auditory scene analysis^37^. Up to now, no repeated stimuli used in stimulus-specific adaptation paradigm exhibited any natural variations leaving unclear the outcome of the present study. Zebra finches intensively repeat their vocalizations with slight variations across renditions. One possible prediction was that natural variations prevented or slowed down changes in auditory responses with stimulus repetition because variants are encoded as distinct stimuli. In such a case, regarding the functional role of the adaptation, the change in adaptation rate could be viewed as maintaining the stimulus detection despite its repetition and beyond that, a focus on individual’s vocalizations. Another outcome would be no influence of variations in the time course of responses because the tolerance of NCM neurons allows them to encode a stimulus as an object regardless acoustic variation. Our results provide support to both predictions. Depending on the time scale, the impact of variations on both responses and the time course of the adaptation differed. This is consistent with studies reporting that cortical auditory neurons exhibiting stimulus-specific adaptation shows a sensitivity to auditory stimuli that operates at multiple time scales concurrently, spanning many orders of magnitude ^38^.

When responses were calculated from firing rates averaged over the entire sequence duration, they showed no clear impact of slight variations in acoustic features of syllables. Responses showed a decrease with stimulus repetition, as described in high-level auditory areas in mammals^39,40^ or songbirds^17,19,20,22,24,41,42^. Importantly, this decrease did not depend on whether variants or same sequences were broadcasted. We reported a similar adaptation rate when greater changes in response magnitude occurred, *i.e*., during the first presentations of the auditory stimuli. This suggests that, at the sequence duration time scale, responsive neurons encode entire sequences as unique objects, independently of the natural acoustic variations of syllables. Consistently, a few studies have previously reported invariance in auditory responses of NCM neurons^13^, even when song stimuli were played back in an environmental background noise^29^. From a temporal perspective, the tolerance of responses to natural acoustic variations does not imply that the length of the time window integrating acoustic information into a single object requires the entire sequence duration. Analysis of temporal patterns of spike trains by varying the Gaussian window width over which convolutions were performed indicated no difference in responses to playbacks of variants and same sequences when time scale exceeded ~100 ms. Consistently, a peak invariance around 150 ms after onset of different call-types has been reported in the avian auditory cortex including the NCM^13^.

Importantly, the present study also provides evidence that, at a short timescale, neuronal responses reflect an impact of the variability in acoustic features of syllables across renditions. Temporal reliability of spike trains was lower when the fine acoustic structure of syllables varied. Also, the time course of the spike train reliability across stimuli differed depending on whether variant or same sequences were played back. The CorrCoef values indeed decreased when the same sequence was used as recurring stimulus while they remained similar when sequences acoustically varied. These results cannot be explained by a lack of temporal organization within spike trains evoked by playbacks of variants that could not allow any decrease in spike timing reliability. Although CorrCoef values were low, they were higher than those for randomly organized spike trains. Also, from the first to the second AB pair within ABAB sequences, CorrCoef values decreased even when sequence variants were used as stimuli. An explanation based on differences in firing rates can also be excluded, the CorrCoef values being independent on the firing rate^26^. Moreover, the firing rate similarly decreased with stimulus repetition even when sequences acoustically varied across renditions. We rather propose that the temporal resolution of spike trains greatly differed depending on whether variants or same sequences were used as stimuli. To compute CorrCoef measures, we performed a convolution of each spike train with a Gaussian window width ranging from 1 to 200 ms. Interestingly, CorrCoef values reached a plateau with a width of about 10 ms when the same sequence was used as repeated stimulus. This implies that the temporal precision of spike trains evoked by similar sequences occurred in a time scale of about 10 ms. In contrast, no clear plateau was reached for spike trains evoked by varying sequences up to 200 ms.

One property of NCM neurons that makes their auditory responses complex is their non-linear integration of acoustic information. The adaptation of responses with stimulus repetition exemplifies this property^20,29,43^. Consistently, we did not find any significant correlation between the temporal patterns of the spike trains acoustic measures (*i.e*., pitch, entropy, similarity score). The lack of a direct contribution of one or a combination of acoustic features in auditory responses of neurons in a high-order brain area may result from a sensitivity to the context in which sound stimuli occurs^20,29,43^. For example, manipulating the temporal order of syllables within songs affected neuronal responses to a given song syllable, neuronal activity depending on which syllable immediately preceded^20^. Here, the repetition of the same AB pair within ABAB sequences offered us the opportunity to examine the impact of global context, variants *vs*. same sequences. The difference observed in temporal patterns of spike trains between the first and the second pair according to the global context provided new support to the idea that neuronal responses in NCM reflect a long-term integration of auditory information that exceeds several hundreds of milliseconds, *i.e*., the time period between the AB pairs of two consecutive sequences. Therefore, NCM neurons were not only sensitive to the fine acoustic structure of syllables, but also to the global context in which syllables occurred. Consistently, such an interplay among multiple time scales in the integration of information was previously described in the auditory cortex of humans^44^ and non-human mammalian species^38,45^ as well as in visual areas^47–49^. Here, a temporal integration scale means the time window during which neurons are sensitive to auditory stimuli, which is different from the time window that can be used to best discriminate between auditory stimuli.

Finally, NCM could provide neural mechanisms to extract critical perceptual information through different types of neural computations based on distinct temporal integration periods: one to provide precise temporal information, one to allow a category to be assigned to the sound stimulus and one to integrate the global context in which sounds occur. These can be related to the richness of behaviorally relevant information encoded in vocal signals, calls and songs^11,46–48^ and to the richness of their temporal structure over multiple time scales^49,50^, as music and speech sounds^51^. A hypothesis based on multiple time integration periods has been proposed for speech and, beyond that, as a general mechanism for audition^44,52,53^.

In summary, our study shows that neurons in a non-primary cortex-like auditory region exhibited sensitivity to fine natural acoustic variations in song elements as well as sensitivity to the context in which song elements occurred, here variants *vs*. similar sequences, suggesting a temporal integration of auditory information across short as well long distinct time scales.

## Methods

### Subjects and housing conditions

The subjects were eleven adult male zebra finches (*Taeniopygia guttata*), reared socially in the breeding colony of the Paris-Saclay University. Birds were kept under a 12:12 light-dark cycle, with food and water *ad libitum*, and an ambient temperature of 22-25°C. Experimental procedures were carried out in compliance with national (JO 887–848) and European (86/609/EEC) legislation on animal experimentation, and following the guidelines used by the animal facilities of Paris-Sud University (Orsay, France), approved by the national directorate of veterinary services (# D91-429).

### Auditory stimuli

Zebra finch song syllables can be categorized into distinct syllable types. To build auditory stimuli, we first selected song syllable types from our collection of song bouts previously recorded (sampling rate: 32 kHz) from adult male zebra finches that had lived in the laboratory’s aviary for years before the experiment. Birds used in the present study had never been exposed to these songs prior to the electrophysiological investigation. A total of 81 syllable types and 60 renditions of each of them were extracted from the bird’s repertoire of twelve male zebra finches. From this dataset, we chose two distinct syllable types, called ‘A’ and ‘B’, that could have been sung by a single or two individuals, to form ABAB sequence stimuli of 0.70 ± 0.30 s duration with 30-50 milliseconds as inter-syllable silence intervals, as typically found in zebra finch songs. Syllable duration ranged from 57 to 235 milliseconds (mean ± SD: 134.2 ± 39.6). Then, we built ABAB-Same series that each consisted of 60 repetitions of a given ABAB sequence (see an example of a ABAB sequence stimulus, called A_1_B_1_A_1_B_1_, in Fig.1) and ABAB-Var series that each consisted of 60 variants of a given ABAB sequence. Variants were labelled as from A_1_B_1_A_1_B_1_ to A_60_B_60_A_60_B_60_ (Fig. 1). Seven ABAB-Same series and eight ABAB-Var series were built. We used Sound Analysis Pro 2011 ^21^ to compute the accuracy score (Fig 1c), which provides a fine-grained quantification of the acoustic similarity, between each renditions of the A and B syllables for each sequences of the ABAB-Same and ABAB-Var series, *i.e*. syllables A *vs* A, B *vs* B, A *vs* B. For the ABAB-Same series for which syllables A and B within a sequence were always the same, an ANOVA revealed a significant difference of the average accuracy scores of the syllables (F_2,28_ = 222.9, *p* < 0.001) and a post-hoc Tukey HSD multiple comparison analysis revealed that it was significantly lower for syllables A *vs* B (average accuracy score = 73.5%) than for syllables A *vs* A (100%) and B *vs* B (100%). For the ABAB-Var series, for which there were 60 variants of the A and B syllables, an ANOVA revealed a significant difference of the average accuracy scores of the syllables (F_2,25_ = 13.93, *p* < 0.001) and a post-hoc Tukey HSD multiple comparison analysis revealed that it was significantly lower for syllables A *vs* B (average accuracy score = 69.2%) than for syllables A *vs* A (82.8%) and B *vs* B (82.4%). None of the ABAB sequences used to build ABAB-Same series were used in ABAB-Var series. All sequences in both series types started with the same introductory note. When a series was played back, sequence stimuli were delivered at a rate of one per second.

### Electrophysiological recordings

Neuronal activity in NCM was recorded in awake (n=4) and in anesthetized (n=7) adult male zebra finches while presenting at least one ABAB-Same and one ABAB-Var series.

### Acute recordings

Birds were anesthetized with isoflurane gas (in oxygen; induction: 3%, maintenance: 1.5%) that flowed through a small mask over the bird’s beak. The bird was immobilized in a custom-made stereotaxic holder that allowed the head to be tilted at 45° and placed in a sound attenuation chamber. Lidocaine cream was applied to the skin. A window was opened in the inner skull layer and small incisions were made in the dura. A multi-electrode array of eight or 16 tungsten electrodes (1-2 MΩ impedance at 1 kHz; Alpha Omega Engineering, Nazareth, Israel) that consisted of two rows of four or eight electrodes separated by 100 μm apart, with 100 μm between electrodes of the same row was lowered to record extracellular activity. The array was positioned 0.3–0.5 mm lateral and 0.7–0.9 mm rostral to the bifurcation of the sagittal sinus in either the left or the right hemisphere, with a micromanipulator, as in previous studies^15,16,20,22^. The probe was lowered very slowly until electrode tips reached 1200 μm below the brain surface. From 1200 to 1900 μm below the brain surface, auditory stimuli were delivered when the amplitude of action potential waveforms recorded with at least one of the eight electrodes was clearly distinct from background noise. Recording sites were at least 100 μm apart to minimize the possibility that the neural activity recorded from two successive sites originated from the same single units. Electrode signals were amplified and filtered (gain 10,000; bandpass: 0.3–10 kHz; AlphaLab SnR, AlphaOmega LTD) to extract multi-unit activity. During recordings, voltage traces and action potentials were monitored in real time using the AlphaLab SnR software. Auditory stimuli were concomitantly recorded and digitized to precisely determine the onset of NCM responses with respect to the sound stimulus. While spiking activity was recorded, auditory stimuli were broadcasted through a loudspeaker situated 30 cm from the bird’s head. We played back one ABAB-Same and one ABAB-Var series. From one recording site to the following one, because of the habituation phenomenon in NCM, we changed the set of series used as auditory stimuli and the order of series. All stimuli had been normalized to achieve maximal amplitude of 70 dB (Audacity software) at the level of the bird’s head. Spike sorting of neuronal activity was done offline (see below).

### Chronic recordings

Surgical procedures were similar as described above. To perform chronic recordings in awake birds, we used a custom build screw microdrive that allows a microelectrode array to be dorsally repositioned. We used arrays of eight electrodes (two rows of four electrodes separated by 100 μm apart; with a ground silver wire and a reference wire; 1-2 MΩ impedance at 1 kHz; Alpha Omega Engineering, Nazareth, Israel). Once the array was lowered into the brain to a depth of 1200 μm, the reference wire was inserted between the outer and the inner skull layers. The microdrive was secured to the skull using dental cement. Subjects were allowed to recover for a few days. In the sound-attenuation chamber, the implanted microdrive was connected through a commercial tether and head stage (AlphaOmega) to a mercury commutator located on the roof of the cage (Dragonfly systems). An elastic thread built into the tether helped to support the weight of the implant. Subjects remained tethered during the experiment. The screw drive held the electrode array. Each full turn of the screw advanced the array by 200 microns. Before a recording session, we rotated the screw by ½ turn to advance the microelectrode array in step as ~100 microns. Birds were not freely moving during the recording session. They were restrained with a jacket around their bodies. At least 24 hours separated two recording sessions. From one recording session to the following one, we changed the set of series used as auditory stimuli.

### Data processing and analysis

In anesthetized birds, spike sorting was performed using the template-matching algorithm of the Spike2 software (version 8.0, Cambridge Electronic Design, CED, Cambridge, UK). NCM contains at least two populations of neurons that can be distinguished on the width of the spike waveform and the firing rate ^20,29,30^, so restricted our analyses to wall-isolated units. In awake birds, neural traces of multiunit activity were subjected to threshold spike detection. Responses to stimuli were quantified by calculating averaged firing rates during sequence presentation and by computing the *RS* index ^15, 22,54^. The *RS* index was calculated by subtracting the spontaneous firing rate (*B_FR_*) from the evoked firing rate (*E_FR_*) and then by dividing this value by their sum:

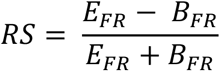

*RS* values fall between +1 and −1, where values >0 indicate an excitatory response and values <0 indicate an inhibitory response. The *B_FR_* was measured over the 200 ms period preceding the stimulus onset. We calculated *RS* values for the 60 renditions of sequence stimuli and per block of 10 presentations, giving us 6 values per series (one per block of ten iterations of the stimulus). Note that for the ABAB-Var series, each block includes 10 variants of the auditory stimuli. Auditory responses to a stimulus in NCM decrease rapidly with stimulus repetition. To examine whether the stimulus-specific adaptation differed between ABAB-Same and -Diff series, we computed a stimulus-specific adaptation rate from responses (*E_FR_*) to the 10 first stimulus renditions by extracting the slope of the linear regression for each unit ^17,23–25^.

The temporal pattern of responses evoked by both types of songs was quantified by calculating the spike-timing reliability coefficient (CorrCoef), which was used to quantify the iteration-to-iteration reliability of responses. It was computed a) per block of ten stimulus iterations and b) per iteration: it corresponds to the normalized covariance between each pair of action potential trains and was calculated as follows:

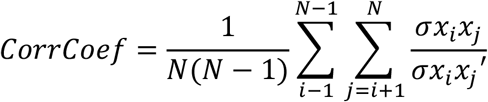

where *N* is the number of iterations, and *xixj* is the normalized covariance at zero lag between spike trains *xi* and *xj*, where *i* and *j* are the iteration numbers. Spike trains *xi* and *xj* were previously convolved with a width of the Gaussian window ranging from 1 to 200 ms. In the present study, most analyses were based on CorrCoef values calculated from a convolution with a 10 ms Gaussian window width, ^20^. The CorrCoef was used because this index is not influenced by fluctuations of firing rate (Gaucher et al, 2013). Note that we also computed CorrCoef values from spikes trains after performing a random permutation of the time at which occurred individual spikes during each stimulus rendition. This random permutation thus gave us an estimation of the CorrCoef when spikes timing is randomly distributed.

Spike-timing reliability might be impacted by the variation of syllables’ duration across each rendition of the ABAB-Var sequences. Given that, we performed a linear time warping of each syllable so that all renditions of an ABAB-Var sequence were aligned on the same time axis ^55^. Syllable boundaries were automatically detected according to the threshold crossing of the root-mean square of the amplitude of each rendition. We extracted the maximum duration of A and B syllables within the sequence and used it as a reference timing. We then linearly stretched or compressed each syllable to match its duration to the maximum duration of its reference. Each individual spike train was then projected to the time warped axis of the corresponding syllable. This algorithm thus reduces the temporal variation of the spike trains from one trial to another.

To examine whether CorrCoef values depended on acoustic variability from one variant to another, we quantified differences in acoustic features and degree of similarity between all variants used to build a given ABAB-Var series with SAP 2011 ^21^. From CorrCoef values computed from spike trains evoked by the two variants used in comparisons, we performed linear regressions.

Statistical computations were carried out in R (4.0.2) and MATLAB (2020a). Firing rates, *RS* and CorrCoef values were analyzed using either repeated measures (RM) ANOVA in Linear Mixed Models (R package ‘nlme’ version 3.1-152) or paired T-tests (R package ‘stats’ version 4.1.0). Depending on the analysis, the block repetition (n=6), the series type (ABAB-Same *vs*. ABAB-Var) and/or AB pair identity (the first *vs*. the second one) were included as cofactors in the model. We used planned contrast and least-square means adjusted with the Tukey HSD tests for assessing pair-wise differences (emmeans function from R package ‘emmeans’ version 1.6.1).

### Histology

At the end of each experiment, the animal was euthanized with a lethal dose of pentobarbital and the brain quickly removed from the skull and placed in a fixative solution (4% para-formaldehyde). Sections (100 μm) were cut on a vibratome to examine the location of multielectrode array penetration tracks.

## Acknowledgements

This work was supported by the Centre National de la Recherche Scientifique, the Idex Neuro-Saclay, and the University of Paris Sud. N.G. was supported by Idex Neuro Saclay Postdoctoral Fellowship. A.C., was supported by the French Ministry of Research and Technology. We thank Chloé Huetz for help in analyzing the data and Jean-Marc Edeline for advices on data interpretation. We thank Mélanie Dumont and Caroline Rousseau for taking care of the songbird facility.

## Author contributions

A.C., N.G., and C.D.N. performed research; A.C., N.G., and C.D.N. analyzed data; N.G. and C.D.N. designed research; N.G. and C.D.N. edited the paper; N.G. wrote the paper.

## Competing interests policy

The authors declare no conflict of interest.

## Data availability

Data will be made available upon reasonable request.

## References

1. Tibbetts, E. A. & Dale, J. Individual recognition: it is good to be different. Trends Ecol Evol 22, 529–537 (2007).

2. Hall, J. A., Horgan, T. G. & Murphy, N. A. Nonverbal communication. Annu Rev Psychol 70, 271–294 (2019).

3. Meliza, C. D. & Margoliash, D. Emergence of selectivity and tolerance in the avian auditory cortex. J Neurosci 32, 15158–15168 (2012).

4. Kanwal, J. S. & Rauschecker, J. P. Auditory cortex of bats and primates: managing species-specific calls for social communication. Front Biosci 12, 4621–4640 (2007).

5. Sharpee, T. O., Nagel, K. I. & Doupe, A. J. Two-dimensional adaptation in the auditory forebrain. J Neurophysiol 106, 1841–1861 (2011).

6. Liu, S. T., Montes-Lourido, P., Wang, X. & Sadagopan, S. Optimal features for auditory categorization. Nat Commun 10, 1302 (2019).

7. Falls, J. B. Individual recognition by sound in birds. in Acoustic communication in birds (eds. Kroodsma, D. E. & Miller, E. H.) vol. 2 237–278 (Academic Press, 1982).

8. Lambrechts, M. M. & Dhondt, A. A. Individual voice discrimination in birds. in Current Ornithology (ed. Power, D. M.) 115–139 (Springer US, 1995).

9. Hyland Bruno, J. & Tchernichovski, O. Regularities in zebra finch song beyond the repeated motif. Behav Proc 163, 53–59 (2019).

10. Woolley, S. C. & Doupe, A. J. Social context-induced song variation affects female behavior and gene expression. PLoS Biol. 6, e62 (2008).

11. Fishbein, A. R., Prior, N. H., Brown, J. A., Ball, G. F. & Dooling, R. J. Discrimination of natural acoustic variation in vocal signals. Sci Rep 11, 916 (2021).

12. Elie, J. E. & Theunissen, F. E. Meaning in the avian auditory cortex: neural representation of communication calls. Eur J Neurosci 41, 546–567 (2015).

13. Elie, J. E. & Theunissen, F. E. Invariant neural responses for sensory categories revealed by the time-varying information for communication calls. PLOS Comput Biol 15, e1006698 (2019).

14. Thompson, J. V. & Gentner, T. Q. Song recognition learning and stimulus-specific weakening of neural responses in the avian auditory forebrain. J Neurophysiol 103, 1785–1797 (2010).

15. Menardy, F. et al. Social experience affects neuronal responses to male calls in adult female zebra finches. Eur J Neurosci 35, 1322–1336 (2012).

16. Menardy, F., Giret, N. & Del Negro, C. The presence of an audience modulates responses to familiar call stimuli in the male zebra finch forebrain. Eur J Neurosci 40, 3338–3350 (2014).

17. Chew, S. J., Mello, C., Nottebohm, F., Jarvis, E. & Vicario, D. S. Decrements in auditory responses to a repeated conspecific song are long-lasting and require two periods of protein synthesis in the songbird forebrain. Proc Natl Acad Sci USA 92, 3406–3410 (1995).

18. Mello, C., Nottebohm, F. & Clayton, D. Repeated exposure to one song leads to a rapid and persistent decline in an immediate early gene’s response to that song in zebra finch telencephalon. J Neurosci 15, 6919–6925 (1995).

19. Beckers, G. J. L. & Gahr, M. Neural processing of short-term recurrence in songbird vocal communication. PLoS ONE 5, e11129 (2010).

20. Cazala, A., Giret, N., Edeline, J.-M. & Del Negro, C. Neuronal encoding in a high-level auditory area: from sequential order of elements to grammatical structure. J Neurosci 39, 6150–6161 (2019).

21. Tchernichovski, O., Nottebohm, F., Ho, C. E., Pesaran, B. & Mitra, P. P. A procedure for an automated measurement of song similarity. Anim Behav 59, 1167–1176 (2000).

22. Stripling, R., Volman, S. F. & Clayton, D. F. Response modulation in the Zebra finch neostriatum: relationship to nuclear gene regulation. J Neurosci 17, 3883–3893 (1997).

23. Chew, S. J., Vicario, D. S. & Nottebohm, F. A large-capacity memory system that recognizes the calls and songs of individual birds. Proc Natl Acad Sci USA 93, 1950–1955 (1996).

24. Phan, M. L., Pytte, C. L. & Vicario, D. S. Early auditory experience generates long-lasting memories that may subserve vocal learning in songbirds. Proc Natl Acad Sci USA 103, 1088–1093 (2006).

25. Terleph, T. A., Mello, C. V. & Vicario, D. S. Auditory topography and temporal response dynamics of canary caudal telencephalon. J Neurobiol 66, 281–292 (2006).

26. Gaucher, Q., Huetz, C., Gourévitch, B. & Edeline, J.-M. Cortical inhibition reduces information redundancy at presentation of communication sounds in the primary auditory cortex. J Neurosci 33, 10713–10728 (2013).

27. Gaucher, Q. & Edeline, J.-M. Stimulus-specific effects of noradrenaline in auditory cortex: implications for the discrimination of communication sounds. J Physiol 593, 1003–1020 (2015).

28. Souffi, S., Lorenzi, C., Varnet, L., Huetz, C. & Edeline, J.-M. Noise-sensitive but more precise subcortical representations coexist with robust cortical encoding of natural vocalizations. J Neurosci 40, 5228–5246 (2020).

29. Schneider, D. M. & Woolley, S. M. N. Sparse and background-invariant coding of vocalizations in auditory scenes. Neuron 79, 141–152 (2013).

30. Ono, S., Okanoya, K. & Seki, Y. Hierarchical emergence of sequence sensitivity in the songbird auditory forebrain. J Comp Physiol A 1–21 (2016) doi:10.1007/s00359-016-1070-7.

31. Yanagihara, S. & Yazaki-Sugiyama, Y. Auditory experience-dependent cortical circuit shaping for memory formation in bird song learning. Nat Commun 7, 11946 (2016).

32. Huetz, C., Del Negro, C., Lebas, N., Tarroux, P. & Edeline, J.-M. Contribution of spike timing to the information transmitted by HVC neurons. Eur J Neurosci 24, 1091–1108 (2006).

33. Narayan, R., Graña, G. & Sen, K. Distinct time scales in cortical discrimination of natural sounds in songbirds. J Neurophysiol 96, 252–258 (2006).

34. Ribeiro, S., Cecchi, G. A., Magnasco, M. O. & Mello, C. V. Toward a song code: evidence for a syllabic representation in the canary brain. Neuron 21, 359–371 (1998).

35. Woolley, S. M. N., Gill, P. R. & Theunissen, F. E. Stimulus-dependent auditory tuning results in synchronous population coding of vocalizations in the songbird midbrain. J. Neurosci. 26, 2499–2512 (2006).

36. Laudanski, J., Edeline, J.-M. & Huetz, C. Differences between spectro-temporal receptive fields derived from artificial and natural stimuli in the auditory cortex. PLOS ONE 7, e50539 (2012).

37. Winkler, I., Denham, S. L. & Nelken, I. Modeling the auditory scene: predictive regularity representations and perceptual objects. Trends Cogn Sci 13, 532–540 (2009).

38. Ulanovsky, N., Las, L., Farkas, D. & Nelken, I. Multiple time scales of adaptation in auditory cortex neurons. J Neurosci 24, 10440–10453 (2004).

39. Malmierca, M. S., Sanchez-Vives, M. V., Escera, C. & Bendixen, A. Neuronal adaptation, novelty detection and regularity encoding in audition. Front Syst Neurosci 8, (2014).

40. Khouri, L. & Nelken, I. Detecting the unexpected. Curr Opin Neurobiol 35, 142–147 (2015).

41. Smulders, T. V. & Jarvis, E. D. Different mechanisms are responsible for dishabituation of electrophysiological auditory responses to a change in acoustic identity than to a change in stimulus location. Neurobiol Learn Mem 106, 163–176 (2013).

42. Lu, K. & Vicario, D. S. Statistical learning of recurring sound patterns encodes auditory objects in songbird forebrain. Proc Natl Acad Sci USA 111, 14553–14558 (2014).

43. Lu, K. & Vicario, D. S. Familiar but unexpected: effects of sound context statistics on auditory responses in the songbird forebrain. J. Neurosci. 37, 12006–12017 (2017).

44. Teng, X., Tian, X. & Poeppel, D. Testing multi-scale processing in the auditory system. Sci Rep 6, 34390 (2016).

45. García-Rosales, F., Beetz, M. J., Cabral-Calderin, Y., Kössl, M. & Hechavarria, J. C. Neuronal coding of multiscale temporal features in communication sequences within the bat auditory cortex. Commun Biol 1, 1–14 (2018).

46. Elie, J. E. & Theunissen, F. E. Zebra finches identify individuals using vocal signatures unique to each call type. Nat Commun 9, 4026 (2018).

47. Perez, E. C. et al. The acoustic expression of stress in a songbird: does corticosterone drive isolation-induced modifications of zebra finch calls? Horm Behav 61, 573–581 (2012).

48. D’Amelio, P. B., Klumb, M., Adreani, M. N., Gahr, M. L. & Maat, A. Individual recognition of opposite sex vocalizations in the zebra finch. Sci Rep 7, 5579 (2017).

49. Cynx, J., Williams, H. & Nottebohm, F. Timbre discrimination in zebra finch (Taeniopygia guttata) song syllables. J Comp Psychol 104, 303–308 (1990).

50. Lohr, B., Dooling, R. J. & Bartone, S. The discrimination of temporal fine structure in call-like harmonic sounds by birds. J Comp Psychol 120, 239–251 (2006).

51. Rosen, S., Carlyon, R. P., Darwin, C. J. & Russell, I. J. Temporal information in speech: acoustic, auditory and linguistic aspects. Philosophical Transactions of the Royal Society of London. Series B: Biological Sciences 336, 367–373 (1992).

52. Poeppel, D. Pure word deafness and the bilateral processing of the speech code. Cogn Sci 25, 679–693 (2001).

53. Poeppel, D. The analysis of speech in different temporal integration windows: cerebral lateralization as ‘asymmetric sampling in time’. Speech Commun 41, 245–255 (2003).

54. Giret, N., Menardy, F. & Del Negro, C. Sex differences in the representation of call stimuli in a songbird secondary auditory area. Front. Behav. Neurosci. 9, 290 (2015).

55. Kao, M. H., Wright, B. D. & Doupe, A. J. Neurons in a forebrain nucleus required for vocal plasticity rapidly switch between precise firing and variable bursting depending on social context. J. Neurosci. 28, 13232–13247 (2008).

